# Seasonal dynamics and thermal vulnerability of marine microbial communities in Uruguayan coastal waters

**DOI:** 10.1101/2025.06.19.660397

**Authors:** Emiliano Pereira, Juan Zanetti, Luciana Griffero, Ana Martínez, Rudolf Amann, Cecilia Alonso

## Abstract

In an era of accelerating ocean warming, understanding the drivers of microbial community composition is more urgent than ever. Long-term time-series studies are particularly powerful for this purpose. However, most previous work remains confined to the Northern Hemisphere and focuses solely on taxonomy, leaving the Southern Hemisphere underrepresented and microorganisms’ functional dimensions underexplored. Moreover, phenomena such as current-driven community shifts and microbial vulnerability to warming remain poorly understood. Here, we address these gaps by analyzing microbial communities over two years at the South Atlantic Microbial Observatory (SAMO), located off the coast of Uruguay within a Marine Protected Area. SAMO, situated under the contrasting influences of the Brazil and Malvinas currents and recognized as a global-warming hotspot, provides an ideal setting to study the temporal dynamics of marine microbial communities. Using shotgun metagenomics and 16S rRNA amplicon sequencing, we revealed strong seasonal shifts driven by environmental variability and the influence of these two ocean currents, resulting in two community states (summer–autumn vs winter–spring). These states exhibited contrasting composition, diversity, and assembly mechanisms. Further, our results show that taxonomic and functional diversity trends diverge between these periods. Crucially, our work suggests that the predicted tropicalization of Uruguay’s coast is likely to boost taxonomic richness but erode functional repertoires, while also increasing vulnerability to ocean warming, potentially undermining biogeochemical functioning.

## Introduction

Marine ecosystems harbor an astonishing microbial diversity and biomass that underpin ecosystem functioning and global biogeochemical cycles (Falkowski 2008). Unraveling the environmental drivers and ecological processes that shape marine microbial communities is fundamental to marine microbial ecology, and all the more vital as rapid global changes, especially those driven by warming, reshape our oceans (Doney 2012). Long-term time-series studies are particularly valuable for characterizing the temporal dynamics and underlying mechanisms that shape the assembly, structure, and diversity of marine microbial communities (Fuhrman 2015). Previous research has shown that these communities exhibit strong seasonal recurrence linked to environmental variability and are largely governed by deterministic processes (Fuhrman 2006; Teeling 2016; Chafee 2018; Ferrera 2024). This recurrence is especially pronounced in euphotic zones at higher latitudes, such as temperate and polar regions (Fuhrman 2015). Overall, the strong and consistent seasonal patterns driven by temperature cycles make long-term time-series studies particularly well suited to assess the impacts of global warming on marine microbial dynamics.

Despite valuable insights of previous studies analysing marine microbial dynamics over time, as those cited above, it is important to note that most of these are from the northern hemisphere, and based only on taxonomic information (i.e., 16S rRNA amplicon data) (Fuhrman 2015; Raes 2024). Currently, key questions remain unanswered. Particularly, it remains unclear whether functional and taxonomic community compositions exhibit comparable patterns of recurrence, diversity, and environmental association; how ocean currents drive shifts in community structure; and how microbial assemblages will respond to rising ocean temperatures. Addressing these gaps is essential for refining biogeochemical models, guiding marine-ecosystem management, and mitigating the consequences of anthropogenic disturbances, most notably, global warming (Cavicchioli 2019; Abirami 2021).

The South Atlantic Microbial Observatory (SAMO), located off the coast of Uruguay, provides a unique opportunity to address these questions. The site is highly influenced by the Brazil and Malvinas currents, which have contrasting environmental characteristics, and is recognized as a global warming hotspot (Ortega 2007; Hobday 2014). This makes the region an ideal natural laboratory to study the dynamics and seasonality of marine microbial communities in the Southern Hemisphere and to assess the influence of ocean currents and climate change on these patterns.

In this study, we analyzed time-series data from SAMO to characterize shifts in coastal microbial communities. The dataset spans two years and includes metagenomic and 16S rRNA amplicon samples, along with environmental descriptors. Our results revealed strong seasonal patterns at both taxonomic and functional levels, driven by environmental variability and the influence of the mentioned currents. Two distinct community types emerged, corresponding to summer/autumn and winter/spring semesters, differing in composition, diversity, assembly mechanisms, and vulnerability to ocean warming. Notably, under the projected tropicalization of Uruguay’s coastal waters (Ortega 2016), microbial communities are expected to show increased taxonomic but reduced functional diversity, along with heightened vulnerability to warming, potentially compromising ecosystem functioning.

## Materials and methods

## 1. Description of the sampling site

Our sampling site, the South Atlantic Microbial Observatory (SAMO), is located two nautical miles off the coast of Rocha, Uruguay (34°42’43.4’’S, 54°14’08.6’’W), within the Laguna de Rocha Marine Protected Area (MPA). It lies adjacent to the Laguna de Rocha and near the Brazil–Malvinas Confluence Zone. SAMO is influenced by warm, oligotrophic water masses advected by the Brazil Current during summer and fall, and by colder, more diluted, nutrient-rich waters from the northward-flowing Malvinas Current in winter and spring. Additional influences include the Río de la Plata, which shows seasonal and interannual variation mainly linked to the El Niño–Southern Oscillation phenomenon, as well as freshwater pulses from the Laguna de Rocha. Phytoplankton blooms occur year-round, with high-biomass dinoflagellate blooms in late winter–early spring and toxic blooms in autumn (Martínez 2019; Martínez 2024).

## 2. Sample collection

### 2.1. Biological material collection

The current study analyzes samples collected at SAMO from a two year period, from 2018-02-20 to 2020-03-11. In total 33 and 37 DNA samples corresponding to the 25-0.22 µm fraction size were subjected to 16S rRNA amplicon and metagenomic (i.e., whole genome shotgun) sequencing, respectively.

### 2.2. Environmental data collection

Physicochemical parameters were measured in situ using a Horiba multiparameter sensor, including conductivity, turbidity, temperature, dissolved oxygen, salinity, total dissolved solids, density, and pH. Chlorophyll and phycocyanin fluorescence were assessed with a Turner fluorometer, and light penetration with a Secchi disc.

## 3. 16S rRNA amplicon sequence data pre-processing, clustering, and taxonomic annotation

From the initial 33 amplicon-sequenced samples, we excluded those with chlorophyll concentrations >7 µg/l, associated with algal blooms, which contained bacterioplankton communities that differed substantially from the rest of the dataset, and obscured environmental and water mass patterns. This filtering yielded 28 samples for 16S rRNA amplicon analysis.

Sequence preprocessing and Amplicon Sequence Variants (ASVs) inference were performed using the DADA2 R package (v1.26.0) (Callahan 2016). Taxonomic annotation was done using the Naive Bayes Classifier (Wang 2007) trained on the Silva NR99 v138 database (Quast 2013). ASVs found in fewer than two samples or classified as mitochondria or chloroplasts were removed, as were samples with fewer than 2,500 reads. To normalize sequencing depth, the abundance table was rarefied to 2,687 sequences per sample, resulting in 26 samples and 3,292 ASVs (Supplementary Table S1 and Supplementary Methods).

## 4. Metagenomic sequence data pre-processing, clustering, and functional annotation

From the 37 metagenomic samples, we selected those matching the amplicon dataset (i.e., same biological sample), with ≥3 million paired-end reads and an average quality score >35.1. This removed a single outlier with low quality and read count, resulting in 22 metagenomes.

Raw metagenomic reads were quality-filtered and trimmed to generate both, preprocessed paired-end and merged reads. OPUs (Operational Protein Units) were constructed by assembling the quality-filtered paired-end reads, predicting Open Reading Frames (ORFs), estimating their coverage, and clustering at 70% amino acid identity. Based on the clustering and coverage estimates, we built OPU abundance tables. We removed OPUs present in fewer than two samples or with total abundance <30, and rarefied the table to a minimum abundance per sample (see Supplementary Tables S2 and S3; and Supplementary Methods).

Functional annotation was performed on OPU representative sequences (pre-rarefaction) by aligning them to the KEGG Ortholog (KO) database (Kanehisa 2017) using hmmsearch from the pyHMMER Python module (Larralde 2023) with a domain e-value threshold of 1e–5.

## 5. Data analysis

### 5.1. Ordination analysis

To visualize temporal microbial beta diversity at taxonomic and functional levels, we performed nonmetric multidimensional scaling (NMDS) using the cmdscale function from the vegan R package v2.6-8 (Oksanen 2024), based on Bray–Curtis dissimilarities of rarefied and Hellinger-transformed ASV and OPU abundance matrices. Samples were grouped by season following the southern hemisphere calendar. Seasonal trajectories were represented by the centroids of each season in the first two NMDS axes.

### 5.2. Time decay analysis

To estimate community time–dissimilarity relationships, we compared the number of days between each sample pair with their Bray–Curtis dissimilarity at taxonomic and functional levels (i.e., ASV and OPU abundance tables). To visualize these associations, we fitted a LOESS curve using the loess.as function from the fANCOVA R package v0.6.1 (Wang 2020), with default parameters.

### 5.3. Space-time analysis

To perform the space-time analysis, we compared microbial communities from SAMO with six TARA Oceans 16S rRNA miTAG samples from the South Atlantic. miTAG sequences from the prokaryote-enriched fraction and their metadata were downloaded from EMBL (http://ocean-microbiome.embl.de) (Sunagawa 2015). Sequences were clustered at 97% identity using VSEARCH v2.15.1 (Rognes 2016) with the cluster_fast method (length-sorted clustering) to generate an OTU abundance table. Centroid sequences were taxonomically annotated using the DADA2 Naive Bayes Classifier, following the same procedure described above. Genus-level abundances (bootstrap >50%) were estimated for both TARA OTUs and SAMO ASVs, and both tables were rarefied to the minimum combined abundance (2,687 reads).

To map SAMO samples, we computed Bray–Curtis dissimilarities at the genus level against six TARA Oceans samples collected from surface waters near the SAMO site—three located to the north and three to the south (see Supplementary Table S4). We then calculated Bray–Curtis–scaled similarities and used them as weights to estimate new geographic coordinates for each SAMO sample (see Supplementary Methods).

### 5.4. Diversity estimations

To estimate taxonomic and functional diversity, we calculated observed richness, the Chao1 richness estimator, and the Shannon diversity index using rarefied ASV and OPU abundance tables. The latter two were computed with the estimateR and diversity functions from the vegan R package.

### 5.5. Loss vs gain comparison

To estimate ASV and OPU gains and losses between semesters, we applied Bray–Curtis dissimilarity decomposition (Legendre 2019). Gains (C) and losses (B) were derived by comparing species abundances, with B representing losses and C gains. We computed B and C across seasonal transitions (Summer–Fall, Winter–Spring, and vice versa), recorded cases where C > B as gains (see Supplementary Methods).

### 5.6. Differential associations and abundance testing

To identify ASVs and KEGG Orthologs (KOs) consistently and significantly associated between semesters, we applied the following approach. First, we generated 100 rarefied ASV and KO abundance tables, each subsampled to the minimum total abundance per sample. For each, we performed the Indicator Value (IndVal) analysis (Dufrêne 1997) using the multipatt function from the indicspecies R package v1.8.0 (De Cáceres 2009), with 9,999 permutations and the seasons Summer-Fall and Winter-Spring as grouping factors. We defined a “consistency” score as the number of times a feature was significant (p ≤ 0.01) across the 100 rarefactions and reported ASVs and KOs with scores of 100 and ≥80, respectively.

In parallel, we used the aldex function from the ALDEx2 R package (Fernandes 2014) to identify differentially abundant ASVs and KOs, considering features significant if both Welch’s t-test and Wilcoxon test yielded Benjamini–Hochberg corrected p-values < 0.05.

### 5.7. Beta Nearest Taxon Index and Nearest Taxon Index

We applied a methodology similar to that described by Logares (2020) to compute the weighted Beta Nearest Taxon Index, and Nearest Taxon Index (bNTI and NTI, respectively) Stegen (2012). Using the most abundant ASVs, we constructed a phylogenetic tree and calculated weighted bNTI and NTI. As required, we tested for phylogenetic signal in environmental preferences over short phylogenetic distances (see Supplementary Methods).

To estimate environmental heterogeneity across semesters (Summer–Fall and Winter–Spring), we followed the approach of Huber (2020), using temperature, salinity, pH, chlorophyll-a, and dissolved oxygen as environmental variables. These variables were scaled and used to compute pairwise Euclidean distances between samples. The resulting distance matrix was normalized by its maximum value and adjusted with a pseudocount (+0.001) to ensure all values were non-zero.

### 5.8. Computation of the 16S rRNA Average Copy Number

To compute the 16S rRNA Average Copy Number (ACN), we applied the run_acn.sh script (Pereira-Flores 2020) to the preprocessed metagenomic samples, consisting of quality-checked merged paired-end reads.

### 5.9. Increase temperature simulations

To assess the vulnerability of microbial communities in each semester under increasing surface water temperature, we conducted a simulation analysis. We first determined the temperature range of each ASV, OPU and semester, then gradually shifted each semester’s range upward by 0 to 3 °C in 0.2 °C increments. ASVs and OPUs with maximum temperature values below the new minimum range were assigned zero abundance. Using the modified abundance tables, we recalculated Shannon diversity and richness (observed and Chao), scaling each metric relative to its original value from the unfiltered table (see Supplementary Methods and Supplementary Figure S1).

### 5.10. Temperature optimum and range analysis

We assessed temperature mismatch in each community by calculating the ratio between each ASV’s optimum and in situ environmental temperatures. Optimum temperature was estimated as the abundance-weighted mean of the temperatures at which each ASV was observed. For each ASV in a sample, we then computed the ratio of its optimum to the in situ temperature. ASV temperature range was defined as the difference between the maximum and minimum temperatures where it was detected. All analyses used the rarefied ASV abundance table.

## Results and Discussion

### Seasonal dynamics

As a first analysis, we explored seasonal dynamics of microbial communities from the Atlantic Uruguayan coast by conducting two NMDS analyses of ASV and OPU beta diversity patterns. The results revealed strong similarities at the taxonomic and functional levels in the seasonal structure of microbial communities and their associations with environmental variability. First, these showed a highly significant differentiation of the semesters of Summer-Fall from Winter-Spring along the first axis (Welch’s ANOVA; all p-values < 0.001). Although the second axis appears to differentiate the semesters of Spring-Summer from Winter-Fall, the trend was much weaker and nonsignificant (Welch’s ANOVA; all p-values > 0.1). At the same time, in both NMDS analyses, we observed a highly significant correlation between the MDS1 and MDS2 axes and the environmental variables temperature and salinity, respectively. While we also observed significant correlations with other environmental variables, temperature and salinity had the highest Pearson correlation values (see Fig. 1A–B; Supplementary Figs. S2–S3; and Supplementary Table S5).

**Figure 1.**
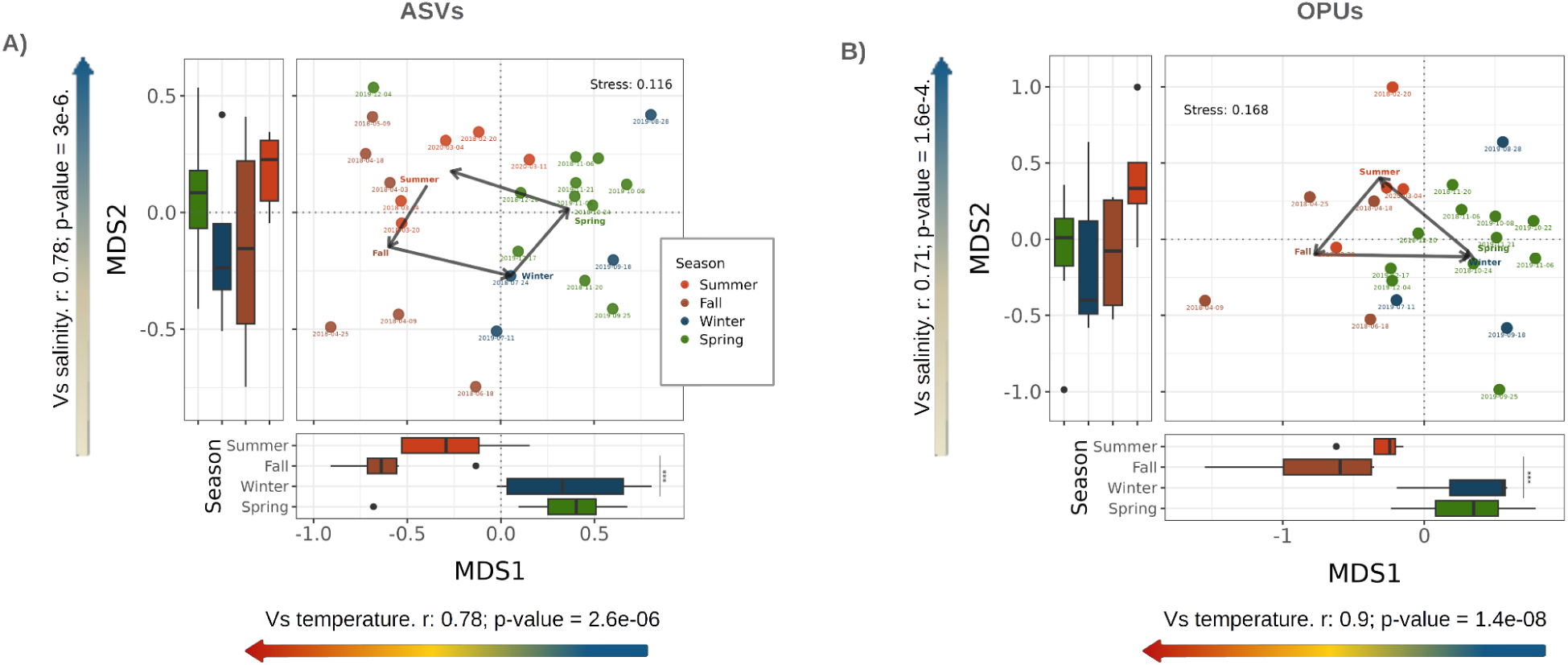
Community differentiation associated with seasonal dynamics and environmental variability. **A and B)** Non-Metric Multidimensional Scaling (NMDS) performed on the Bray-Curtis compositional dissimilarity matrices of Amplicon Sequence Variants (ASVs) and Operational Protein Units (OPUs) abundance profiles. Samples are color coded to represent different seasons. The arrows in the scatter plot connect the centroid of each season showing the transitions between communities over time. The boxplots in the bottom and left sections of the panel show the MDS1 and MDS2 value distributions, respectively, of the samples obtained in the different seasons. In both NMDS analyses, the MDS1 axis differentiated Summer and Fall from Winter and Spring (ANOVA Welch test; all p-values < 0.001). The colored arrows parallel to the MDS1 and MDS2 axes, in the bottom and left sections of the panel, represent the environmental variables temperature and salinity, respectively. In both NMDS analysis, we observed a significant correlation between MDS1 and temperature (ASV composition: pearsons’ r = 0.78 and a p-value < 1e-5; OPU composition: pearsons’ r = 0.9 and a p-value < 1e-7), and MDS2 and salinity (ASV composition: pearsons’ r = 0.78 and a p-value < 1e-5; OPU composition: pearsons’ r = 0.9 and a p-value < 1e-7).

This exploratory analysis indicates that microbial communities are strongly influenced by physical and chemical factors that occur on a seasonal basis, resulting in the differentiation of two main community types. These results resemble a times series study from Ward 2017, performed in a temperate coastal marine ecosystem at the Pivers Island Coastal Observatory (PICO), having practically the same latitude magnitude but opposite sign as ours (i.e., 34.7° N and 34.7 °S, PICO and SAMO, respectively). Namely, as reported by these authors, microbial communities are partitioned into Summer and Winter groups, showing also a strong association with temperature and salinity variability. Although temperature is often reported as the main driver of microbial communities (Sunagawa 2015), the case of salinity is not so obvious, as it can vary depending on the environmental gradient under study (Lozupone 2007).

The results also implied a recurrence of the microbial communities. That is, microbial communities sampled from the same season tend to have a greater similarity, irrespective of the year in which these were sampled. Such recurring patterns can be seen in a trajectory analysis included in the scatterplots of Fig 1 A and B, where we represent the transition between the centroid of each season in the first plane of the NMDS. Moreover, when performing a time-decay analysis comparing the Bray-Curtis dissimilarity and time difference between every pair of samples, we observed a notorious increase and decrease in dissimilarity at approximately 180 and 360 days, respectively (see Supplementary Fig. 4). Thus, our work further confirms the recurrence patterns of microbial communities from surface marine environments at the taxonomic and functional levels, in the southern hemisphere, as previously reported (e.g., Fuhrman 2006; Galand 2018; Priest 2025). Such recurrences also suggest that deterministic processes have a relevant role controlling the composition and structure of microbial communities, as discussed below.

### Time-space analysis

Given that SAMO is highly influenced by the Malvinas and Brazil ocean currents (MC and BC, respectively) throughout the year (Ortega 2007), it provides an ideal setting to assess their impact on microbial dynamics. We therefore conducted a space–time analysis comparing our amplicon samples with six miTag TARA Ocean samples from north and south of the SAMO station (see Supplementary Table S4). We assigned a new set of latitude and longitude coordinates to our amplicon samples representing their similarity to the six TARA Ocean samples. That is, samples that were more similar to TARA samples from the north (or south) were moved to the north (or south) accordingly (see methods section Space-time analysis). Upon analysis, we observed that Summer–Fall samples are shifted to the north, indicating a greater influence of the BC, whereas Winter–Spring samples are shifted to the south, indicating a greater influence of the MC (see Figure 2).

**Figure 2.**
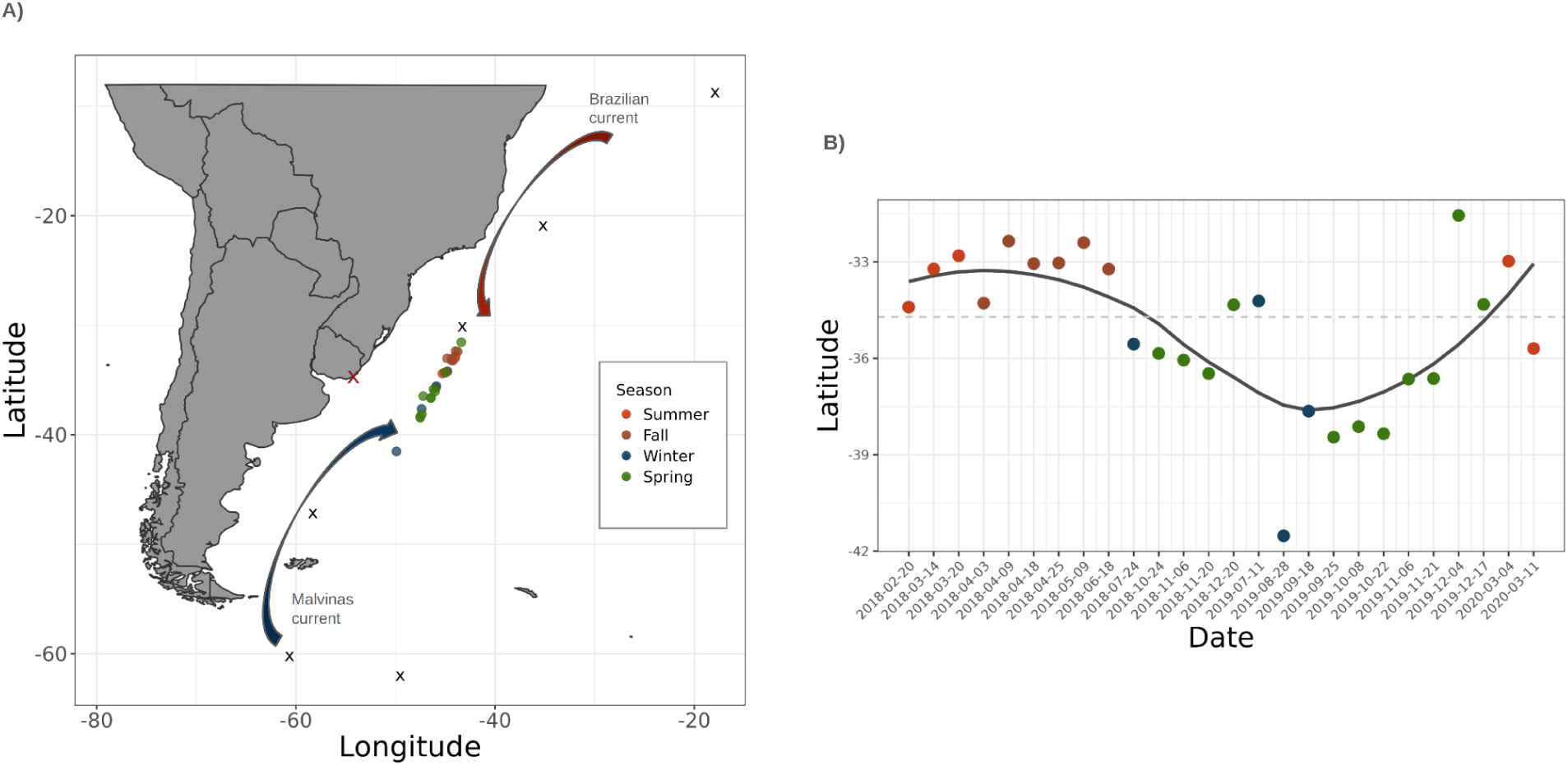
Community differentiation associated with ocean currents dynamics. **A)** Space-time analysis comparing the taxonomic composition of the SAMO and TARA Oceans datasets at the genus level. To position the SAMO samples on the map, they were compared against six TARA Ocean samples from the Southern Atlantic using the Bray-Curtis similarity of genus abundance profiles. The geographic coordinates for each SAMO sample were then calculated as the weighted average of the TARA Ocean samples’ coordinates, with the weights determined by their Bray-Curtis similarity. Black crosses represent the location of the six TARA Ocean samples, and the red cross the location of SAMO. Samples are color coded to represent different seasons. For illustration purposes, the plot shows a schematic representation of the Brazil and Malvinas ocean currents. **B)** Scatterplot showing the variability of the imputed latitude coordinate (as described above) over time. The fitted line was computed utilizing a LOESS regression.

Compared with the MC, the BC is characterized by a higher temperature, salinity and lower nutrient concentrations. While the BC has a dominant presence in the coast of Uruguay during Summer and Fall, MC dominates during Winter and Spring. Here, we show that the recurrence pattern of microbial communities at SAMO, and particularly the strong differentiation of microbial communities between Summer-Fall vs Winter-Spring appears to reflect the influence of these two currents, in agreement with their respective environmental characteristics (Ortega 2007).

The compositional difference between communities from different semesters, and how these in turn reflect the influence of different ocean currents, is congruent with the known ecological differentiation starting around 40°S latitude, where a sharp transition in microbial community composition has been observed (Salazar 2019).

It is highly relevant to relate these results with the previously reported long-term poleward shift of the BC, increasingly linearly with time at approximately 9 km per year (Ortega 2016). This phenomenon, associated with ocean global warming, gives greater significance to our results. It implies that the Summer-Fall communities, which showed a greater similarity with communities from the coast of Brazil, will have a greater dominance in the coast of Uruguay and reach higher latitudes as this trend intensifies.

### Contrasting diversity patterns between semesters

The strong differentiation of microbial communities from the semesters Summer-Fall vs Winter-Spring, and their association with two major ocean currents, prompted us to compare the main community descriptors from each semester. That is, we compared the richness (Chao1 estimator and observed) and the Shannon diversity between semesters at the taxonomic and functional levels. This analysis revealed opposite diversity patterns between semesters. While Summer-Fall had a greater taxonomic diversity and lower functional diversity, Winter-Spring had a lower taxonomic diversity and greater functional diversity. We note that, although we see a consistent pattern, we only obtained a significant difference for the taxonomic richness (i.e., Chao1 estimator and observed; Welch’s ANOVA; all p-values < 0.05).

The greater taxonomic richness during Summer-Fall, is consistent with the fact that this period has greater influence on the BC, transporting warmer and lower-latitude and lower-nutrient waters. Such environmental characteristics are known to contain higher species richness (Ibarbalz 2019). Further, a greater diversity in Summer has been previously reported in either the free living and attached communities (Ward 2017; Mestre 2020) in time series studies; however, this results it is at odds with previous works reporting a higher species richness in Winter (Raes 2024). Clearly, the relationship between seasonality and microbial diversity is complex and may vary depending on specific environmental contexts, such as influence of ocean currents, upwellings, and mixing of water masses.

We found no significant differences between the functional diversity indices; nonetheless, our results suggest that Winter–Spring communities exhibit comparable, and possibly greater, richness than those in Summer–Fall (see Fig. 3B). These results also indicate that microbial communities from Winter–Spring tend to exhibit a greater number of distinct functions per ASV, possibly resulting in lower functional redundancy. Although there are time series studies of marine microbial communities which include functional information obtained from metagenomic data (e.g., Yoshitake 2021; Galand 2018; Priest 2025), these are much scarce, and to our knowledge none of them analyze how the functional alpha diversity varies across the year. Overall, this analysis revealed that taxonomy and function are not necessarily coupled and underscores the need for functional information to link microbial dynamics to ecosystem processes.

**Figure 3.**
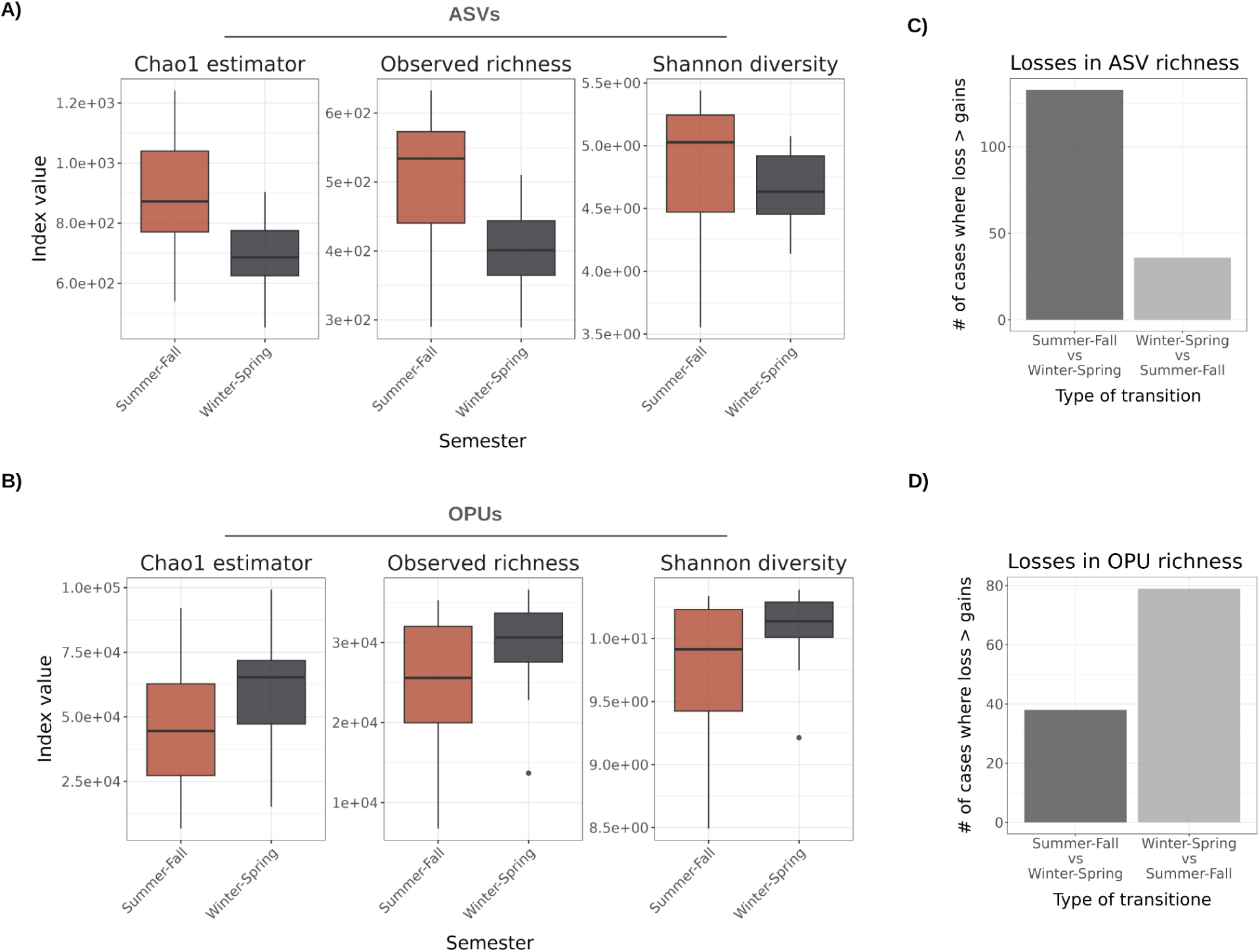
Contrasting diversity patterns between semesters. **A and B)** Box plots comparing the Chao1 estimator, observed richness and Shannon diversity index between semesters Summer-Fall and Winter-Spring at the taxonomic (i.e., ASV composition) and functional (OPU composition) levels, respectively. Significantly different distributions are marked with an asterisk (Welch ANOVA test; all p-values < 0.05). All diversity estimates were computed on the rarefied ASV and OPU abundance matrices. **C and D)** Barplot representing the cases in which the dissimilarity between communities from the Summer-Fall and Winter-Spring is dominated by losses ASVs and OPUs, respectively. We computed the Bray Curtis dissimilarity between all pairs of microbial communities that originate from different semesters (i.e., Summer-Fall vs Winter-Spring), and decomposed it into the loss and gain components as performed in Legendre et al 2019. Next, we counted all cases in which the Bray Curstis dissimilarity was dominated by loss of richness (i.e., the loss component was greater than the gain component). Note that the loss and gain components obtained from the comparison of Summer-Fall vs Winter-Spring are inverted when doing the comparison of Winter-Spring vs Summer-Fall. That is, all the cases in which the loss is greater than the gain in one comparison, will be inverted when the comparison is inverted.

To explore how such contrasting patterns impact the losses and gains of diversity when transitioning between semesters, we applied a decomposition of the community dissimilarities, similarly as implemented in the temporal beta-diversity Index (TBI) of Legendre 2019. In essence, such an approach decomposes the compositional difference, in our case measured by the Bray-Curtis dissimilarity on presence/absence data (i.e., Sorensen dissimilarity), into the gain and loss components (see methods section 5.5). This analysis allows us to explicitly show how the transitions from high to low taxonomically (or functionally) rich communities result in a loss of operational taxonomic (or functional) units. We found that semester-to-semester shifts involved losses of taxonomic groups when moving from Summer–Fall to Winter–Spring, and functional groups in the opposite transition, in spite of the fact that a significant difference was only detected for the taxonomic richness (figs. 3 C and D).

### Differentially associated taxa and functions between semesters

To further investigate this contrasting semester pattern, we applied the Indicator Value (IndVal) analysis (Dufrêne 1997) to the ASV and KEGG Ortholog (KO) (Kanehisa 2017) community compositions to identify the taxonomic groups and functions associated with each semester. In Figure 4 we show the relative abundance of highly consistent and significantly associated ASVs (grouped and color by family and phylum affiliation, respectively) and KOs (see Supplementary File S1 for an extended list of ASVs and KOs significantly associated at consistency scores ≥ 50)).

**Figure 4.**
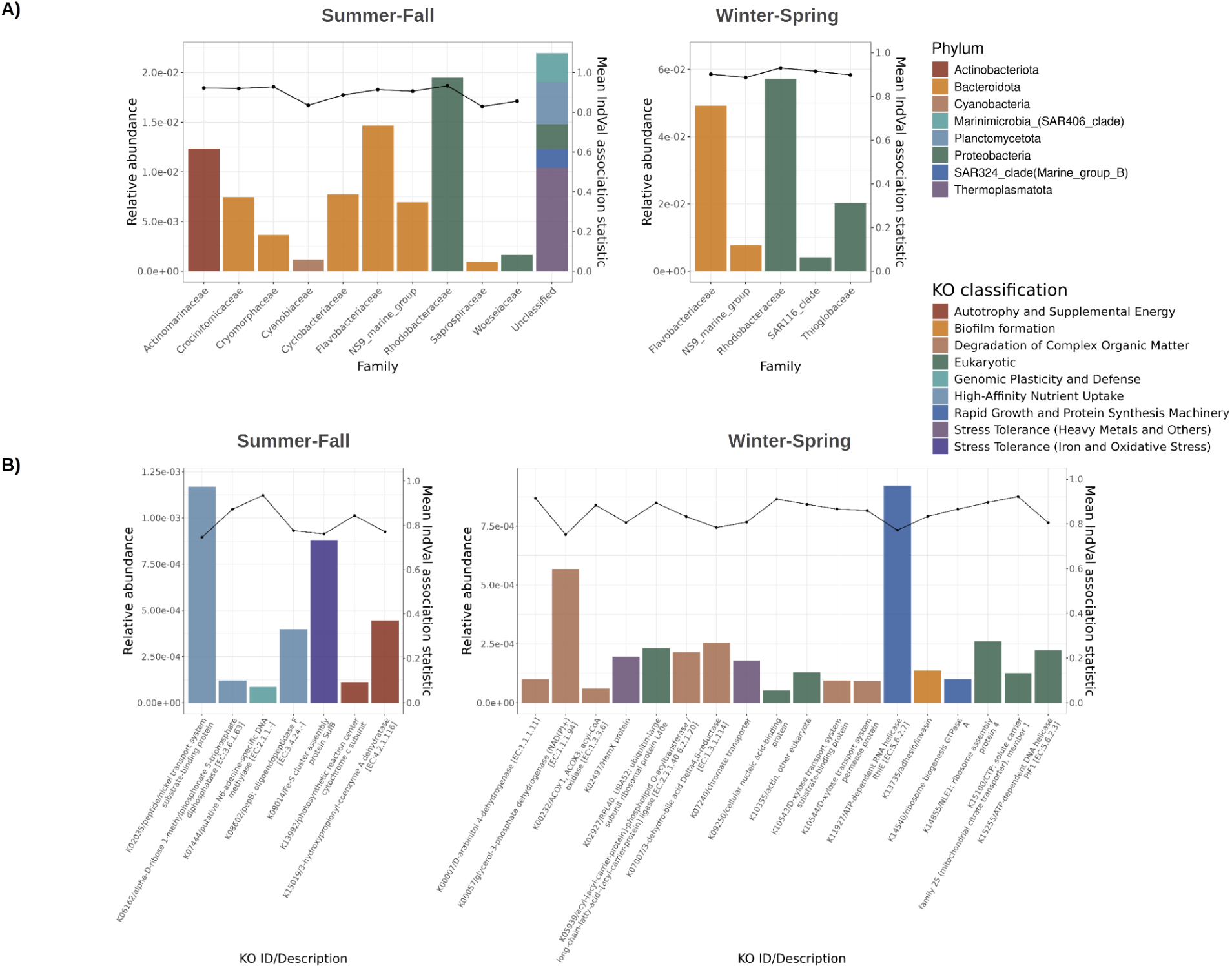
Differentially associated taxa and functions between semesters. **(A)** Bar plot showing the summed relative abundance of ASVs, grouped by family affiliation, that are significantly associated with the Summer–Fall and Winter–Spring semesters (left and right panels, respectively), using the indicator value (IndVal) approach (Dufrêne 1997). Significant ASVs were those with p < 0.01 in all 100 rarefactions of the ASV abundance matrix. The right y-axis represents each family’s mean IndVal association statistic, averaged across all rarefactions and ASVs within that family. Bar colors indicate phylum affiliation. The “Unclassified” column corresponds to ASVs lacking family-level taxonomic annotation. **(B)** Bar plot showing the relative abundance of KEGG Orthologs (KOs) that are significantly associated with the Summer–Fall and Winter–Spring semesters (left and right panels, respectively), also using IndVal. Significant KOs were those with p < 0.01 in at least 80 of the 100 rarefactions of the KO abundance matrix. The right y-axis represents each KO’s mean IndVal association statistic, averaged across all rarefactions.

In line with the previous diversity analysis, the semester Summer-Fall showed a greater number of ASVs and taxonomic groups compared with Winter-Spring (37 vs 11 ASVs, resp.). In the case of the KOs, we observed an opposite pattern: fewer KOs are significantly associated with Summer-Fall vs Winter-Spring (7 and 18 KOs, resp.). These results further suggest that there is a loss of taxonomic diversity when transitioning from Summer-Fall to Winter-Spring and of functional diversity when transitioning from Winter-Spring to Summer-Fall.

In agreement with previous studies, members of the phyla Bateroidota and Proteobacteria, including several families such as NS9_marine_group, Rhodobacteraceae, Flavobacteriaceae, showed a seasonal patterns (Gilbert 2012; Lambert 2021; Ferrera 2024). Also, the latter two families dominated the relative abundance in both semesters (computed as the summed relative abundance of all ASVs within each family). Further, as documented by Ward (2017), our analysis revealed the presence of different ecotypes, indicating temporal niche partitioning (i.e., the NS2b_marine_group and NS4_marine_group genera, from the Flavobacteriaceae family, had representative ASVs in both semesters).

Interestingly, these results showed hallmarks of more oligotrophic and copiotrophic influenced conditions during Summer-Fall and Winter-Spring, respectively, consistent with our findings relating these seasons with the MC and BC. For example, Summer-Fall included significantly associated taxa known to be or in agreement with being highly efficient, light-enhanced, and metabolically streamlined, such as SAR86 bacteria (Roda-Garcia 2023), Candidatus Actinomarinales (López-Pérez 2020), and Marine_Group_II (Zhang 2015). At the functional level, we also see adaptations to nutrient scarcity (K02035, K08602, and K06162) and autotrophy or supplemental energy related functions (K15019 and K13992).

During Winter-Spring, significantly associated taxa appear to be more metabolically versatile, and in agreement with a greater availability of nutrients and resources derived from phytoplankton blooms. An example of this are the genera Aurantivirga, Formosa, and Amylibacter (Sidhu 2023). This observation is echoed at the functional level, where there we find a dominance of functions related with the degradation of complex organic matter (e.g., K10543, K10544, and K05939), rapid growth machinery (K11927 and K14540), and Eucarytic genes (e.g., K02927, K09250, and K15100).

We acknowledge that the alignment of oligotrophic and copiotrophic communities across semesters represents only general tendencies. Although our results reveal clear differentiation between semesters, in reality communities exist along a continuum of states. Moreover, the coastal setting implies year-round influence from continental runoff and nutrient inputs.

Lastly, we note that, although all IndVal association statistics were relatively high, indicating that the detected ASVs and KOs are present in the majority of the samples of each semester, and only in those samples (Dufrêne 1997), the KOs tend to have somewhat lower values. In line with this finding, we observed that the majority of the differentially associated ASVs, genera, and families (62.5, 81.3 and 86.7%, respectively) were also detected as differentially abundant with ALDEX2 (Fernandes 2014). However, ALDEX2 was not able to detect any differentially abundant KOs. This appears to be a consequence of the fact that we are working with relatively shallow metagenomics samples, which in absence of particularly strong signals, between-group differences were too small compared to within-group variability.

### Characterizing and contrasting microbial ecological process

As previously mentioned, the recurrence of microbial communities and their strong association with environmental variables suggest that deterministic processes play significant roles in community assembly. Such observation is further confirmed in the community assembly analysis performed in this work, based on the Nearest Taxon Index (NTI) and beta NTI (bNTI). NTI quantifies the degree of phylogenetic clustering or overdispersion in a community, while bTNI the degree of phylogenetic turnover between two communities (see Materials and Methods section 5.7) (Stegen 2012).

The results showed that the medians of the NTI and bNTI values obtained in both semesters are significantly greater than zero (Wilcoxon Signed Rank Sum test, all p-values < 0.05). These indicate that environmental filtering and heterogeneous selection (i.e., more phylogenetically related and greater phylogenetic turnover than what would be expected under the null model, respectively) have significant roles governing community assembly. Such observation is in line with previous works, evidencing the presence of deterministic processes on prokaryotic microbial communities from marine surface environments (Furhman 2006, Aalto 2022, Priest 2025). Such deterministic processes suggest that shifts in microbial community composition and associated ecosystem functions in response to environmental changes are, to some extent, predictable.

Interestingly, we also observed that the influence of environmental filtering and heterogeneous selection varies for the different semesters. Compared with Summer-Fall, Winter-Spring had a significantly greater mean NTI (Welch’s ANOVA; p-value < 0.05) and a greater number of samples with an NTI value significantly greater than zero (NTI > 2; significantly different from zero at 5% error rate), when analyzed individually (33 vs 93%, Summer-Fall and Winter-Spring, respectively). Considering our previous results, we hypothesize that resource rich environments, as the case of Winter-Spring water masses with a greater influence of the MC, could impose a selection for metabolic capabilities adapted to the local resource characteristics. Conversely, under warmer, more oligotrophic environmental conditions, as in Summer–Fall water masses with greater Brazil Current influence, selection for such metabolic capabilities would be less pronounced. However, in this environment, adaptations to more oligotrophic conditions (resembling K-selected), should be favored (Abreu 2023). In line with this hypothesis, we observed a significant correlation between average 16S rRNA gene copy number and temperature in our dataset (Pearson’s r = 0.45, p = 0.035; see Supplementary Figure S5). Warmer waters tend to harbor a greater number of slow-growing taxa compared with colder waters. Considering that the metabolic profiles should be more susceptible to selection over short-evolutionary time-scales, compared with oligotrophic-related adaptations (Martiny 2013), it would be expected to detect a greater influence of environmental filtering when using nearest taxon indices in Winter-Spring (Stegen 2012).

On the other hand, in the case of the bNTI, we observed a pattern somewhat opposed to that of the NTI: Summer-Fall tends to have greater bNTI values (although the difference was not significant: Welch’s ANOVA; p-value = 0.28) and a greater number of significantly over dispersed community transitions (i.e., bNTI > 2) (55 vs 23%, Summer-Fall and Winter-Spring, respectively). This result is in agreement with the greater environmental heterogeneity captured in the samples from this semester (see Supplementary Figure S6). Further, the fact that environmental filtering has reduced impact during this semester, could be giving more relevance to heterogeneous selection process driven by biotic interactions.

As required, prior to the computation of the NTI and bNTI, we showed that ASVs phylogenetic and environmental distances are significantly correlated when restricted to relatively short phylogenetic distances (see Supplementary Figure S7).

### Increase temperature simulations

SAMO’s lies within a region in which two major ocean currents collide, making it a valuable location for studying climate change potential impact on two contrasting marine environmental conditions. Additionally, the fact that microbial communities show strong deterministic signals, particularly environmental filtering, underscores the utility of this data for forecasting ecological responses to environmental perturbations. With such considerations in mind, we performed a simulation analysis to explore the impact that increasing temperature scenarios can have on the microbial communicates from the different semesters. This analysis consisted of determining the temperature range of each ASV, OPU, and semester, to then artificially shift each semesters’ range towards higher values by small increments, removing all ASVs and OPUs for which their range did not overlap with the semesters’ range (see Materials and Methods section 5.10, and Supplementary Figure S1).

The results show that communities from the Semester Summer-Fall are noticeably more prone to diversity loss in all scenarios of increased temperature at both the taxonomic and functional levels (see Figure 6). The ASV and OPU diversity and richness differences between semesters, became significant (ANOVA Welch test; p-values < 0.05) at increments equal or above 1 and 1.2 degrees Celisius, respectively. For example, the mean Chao1 index computed on the ASV composition dropped to 81% of its initial value after a 1 degree Celsius temperature increment in the Summer-Fall associated communities. Conversely, the same increment only reduced this index to 98% of its initial value in the Winter-Spring associated communities. These results are in line with previous work indicating a greater susceptibility and micro and macro diversity loss caused by global warming in marine tropical environments (Thomas 2012; Ibarbalz 2019; Trisos 2020; Hao 2023).

**Figure 5.**
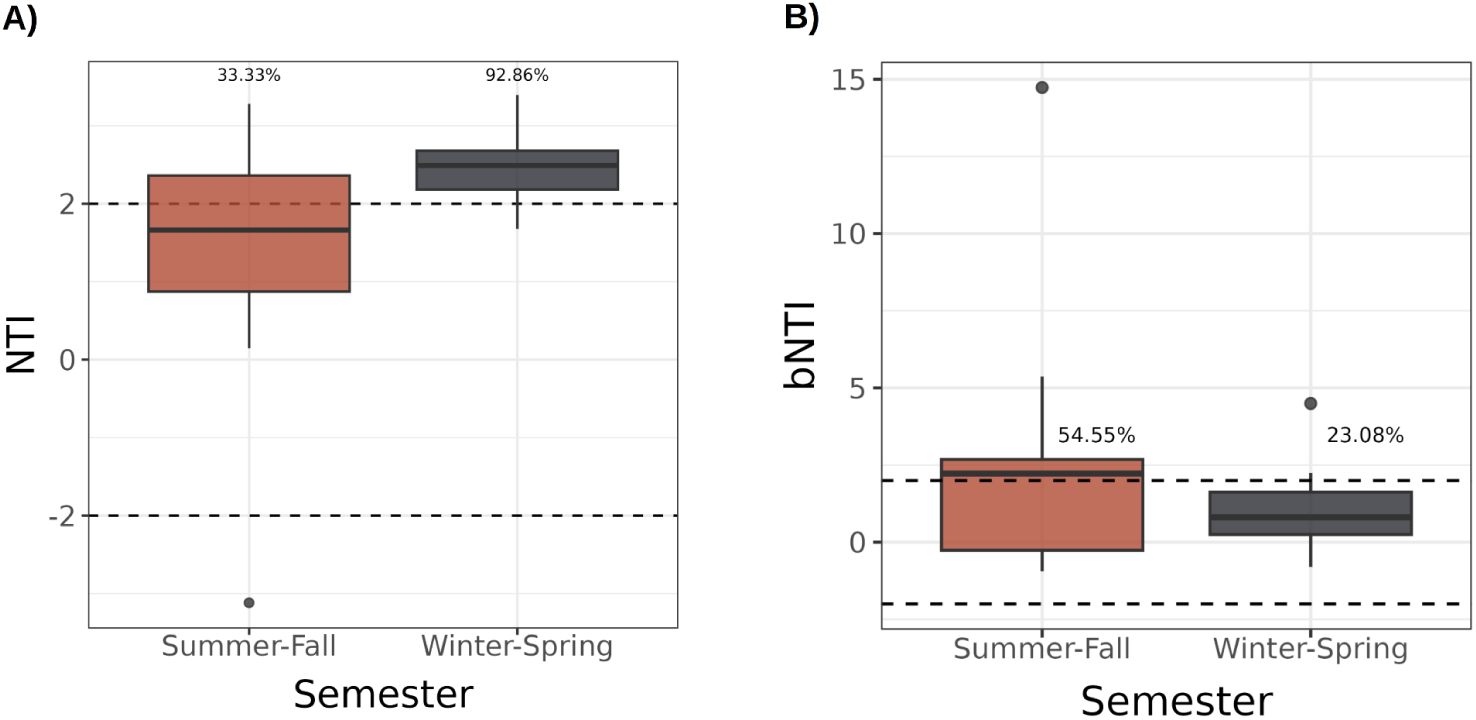
Ecological processes shaping community composition and assembly. A and B) Boxplots showing the distribution of the Nearest Taxon Index (NTI) and beta Nearest Taxon Index (bNTI) values for samples from the Summer-Fall and Winter-Spring semesters, respectively. The median of the NTI and bNTI distributions for both semesters was significantly greater than zero (Wilcoxon Signed Rank Sum test; all p-values < 0.05). Further, the semester Winter-Spring showed significantly greater NTI values compared to the Summer-Fall (Welch’s ANOVA, p-value = 0.035). On the contrary, we did not detect a significant difference of the bNTI distribution between semesters (Welch’s ANOVA, p-value = 0.28). The percentage values above the boxes represent the proportion of samples with an NTI value greater than 2.

**Figure 6.**
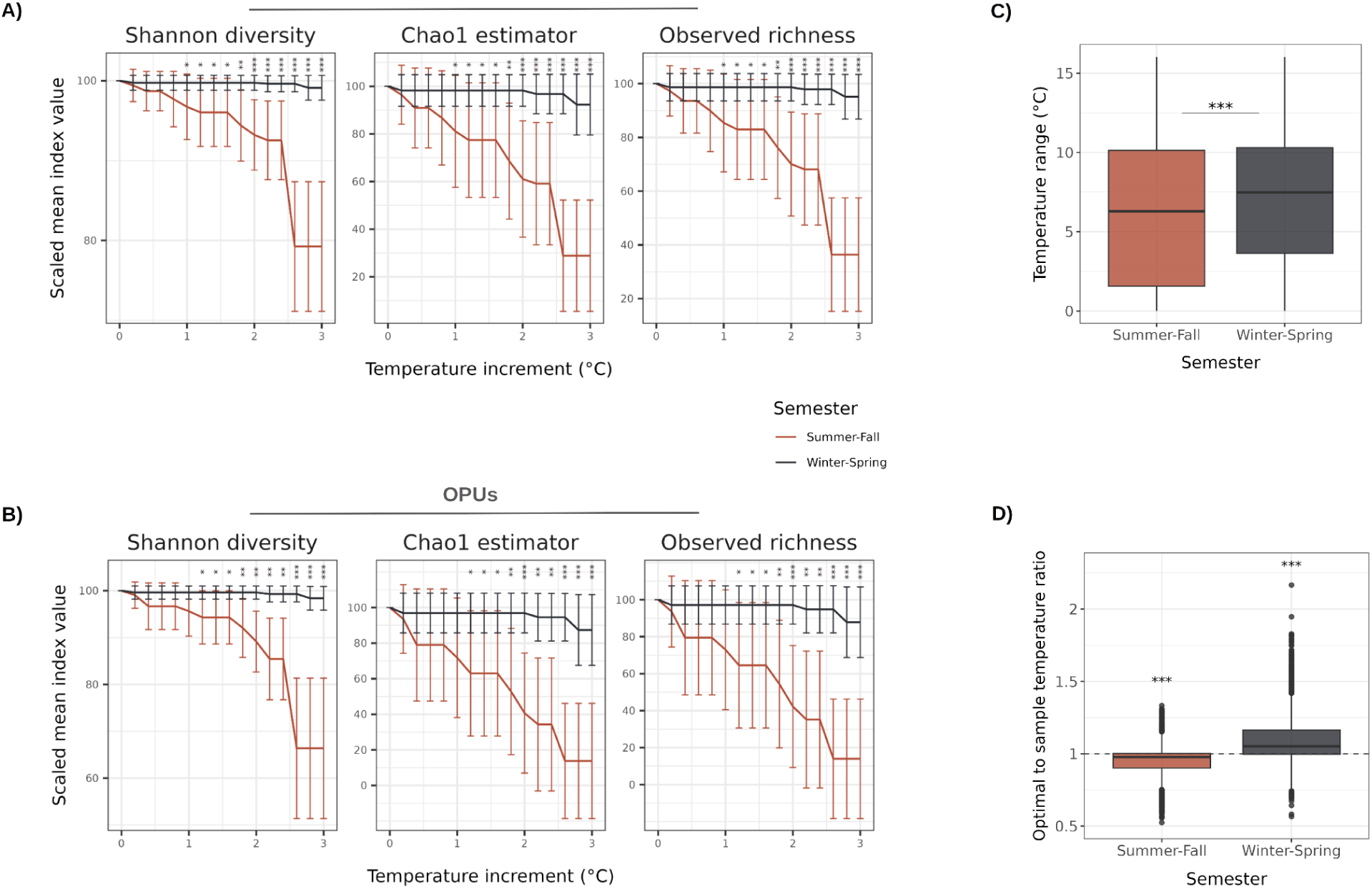
Simulation of increased water temperature scenarios. Figures A and B, illustrate simulated scenarios of temperature increments from 0 to 3 degree Celsius in steps of 0.2, on the ASV and OPU community composition, respectively. Each increment simulation for each semester consisted of removing the ASVs and OPUs for which their range’s temperature maximum was lower than the semester’s range minimum plus the increment. After each temperature increment, three diversity indices (i.e, Shannon diversity, Chao1 index, and Observed richness) were recomputed on the ASVs and OPUs compositions, and scaled by the original values, that is, before any ASVs or OTUs removal. The y-axis represents scaled mean index values and the x-axis the temperature increments (°C). Whiskers represent the first standard deviation across the different samples in each semester. The Summer-Fall and Winter-Spring color-coded as red and blue, respectively. Differences between semesters were tested with ANOVA Welch test and the significance cases are denoted by symbols in the upper section of the plots (i.e., p values lower than 0.05, 0.01, 0.001, represented as one, two and three asterisks, respectively. C) Boxplots representing the temperature range (calculated as the maximum minus the minimum temperature in which an ASV is found) of ASVs from semesters Summer–Fall and Winter–Spring. ASVs from Winter-Spring have an average temperature range significantly greater than Summer-Fall (Welch’s ANOVA, p-value < 2.2e-16). D) Boxplots representing the distribution of the ratio between ASV’s optimal temperature (calculated as the abundance-weighted mean temperature) and the in situ temperature of the sample in which it was detected, for the semesters Summer–Fall and Winter–Spring. A ratio of 1 (indicated by the dashed horizontal line) represents perfect thermal alignment between ASV preference and environmental temperature. Summer-Fall and Winter–Spring show optimal to sample temperature ratio significantly lower and greater than 1 (Wilcoxon Rank Sum Test, p-value < 0.001), as indicated by asterisks above each boxplot.

To evaluate the robustness of our results, we performed the same simulation analysis excluding doubletons and tripletons, or ASVs with an occupancy lower than two, which yield very similar results (data not shown). Further, we demonstrate that the abundance of removed ASVs and OPUs is not significantly different between semesters (Supplementary Figure S8) and includes highly abundant variants, mainly in the case of the communities from Summer-Fall. These control analyses indicate that the pattern is, at least, not solely driven by rare taxa.

Additionally, we compared the temperature range and optimal-to-sample temperature ratio between semesters. In the former case, we observed significant lower values in Summer-Fall (Welch’s ANOVA tests, all p-values < 2.2e-16) compared with Winter-Spring. In the latter case, we observed that Summer-Fall and Winter-Spring have an optimal-to-sample temperature ratio significantly lower and greater than one, respectively (Wilcoxon Rank Sum Test, p-value < 0.001). These results provide an explanation for the greater vulnerability to increasing temperatures observed in the microbial communities from Summer-Fall. First, these results are in agreement with more stable environmental conditions known to occur in tropical marine environments, which as shown in this work, have a greater influence during Summer-Fall. Also, similar observations were made by Chaffron 2021, showing narrower temperature tolerance ranges in tropical and polar associated environments, compared with temperate. Secondly, these results extend the findings of Thomas (2012), which were originally based on a subset of phytoplankton species, to prokaryotic microbial communities in the 25–0.22 µm size fraction. Consistent with the patterns reported by these authors, we show that communities from temperate and tropical environments tend to have optimal growth temperatures that are higher and lower than the prevailing environmental temperatures, respectively. In the latter case, corresponding to Summer-Fall communities, this suggests that rising environmental temperatures will exacerbate the thermal mismatch with species’ optima, potentially increasing physiological stress and altering community dynamics.

Although we observed robust patterns regarding the vulnerability of these Summer-Fall communities to environmental changes, we note that this is an oversimplified simulation. For example, this is based on relatively small datasets, we are not considering other variables that co-vary with temperature (eg., OD), or the immigration of other taxa that could find these environmental conditions suitable, or even evolve (Abirami 2021).

## Conclusions

Marine microbial communities off the Uruguayan coast form two recurring, seasonally distinct assemblages (i.e., Summer–Fall and Winter–Spring) shaped by deterministic processes. Their temporal dynamics closely track environmental shifts in temperature and salinity and the influences of the Brazil and Malvinas Currents, resulting in communities more adapted to oligotrophic conditions during Summer–Fall and copiotrophic conditions during Winter–Spring.

Under the expected tropicalization of the Oceans, our results indicate that these communities will increasingly show a greater taxonomic diversity and lower functional diversity. However, these also appear to be more susceptible to global warming, potentially reducing taxonomic and functional diversity, and in turn, triggering disruptions at ecosystem level.

Considering that SAMO falls within a warming hotspot, we hope that the data generated and analyses performed in this study serve as a baseline for better forecasting and exploring the effects of the warming of the global oceans.

