## Supplementary material for "Seasonal dynamics and thermal vulnerability of marine microbial communities in Uruguayan coastal waters"

### Supplementary methods

#### 16S rRNA amplicon sequence data preprocessing and clustering

To preprocess the 16S rRNA amplicon sequences, we first removed primers using Cutadapt v4.4 (Martin 2011). We then applied the DADA2 R package v1.26.0 (Callahan 2016) following this workflow: 1) Quality trimming and filtering using the filterAndTrim function with the truncLen parameter set to c(220, 175) (i.e., R1 and R2 reads trimmed to 220 and 175 bp, respectively); 2) Dereplication of reads; 3) Merging of paired-end reads with a minimum overlap of 12 bp; 4) Chimera removal; 5) Inference of Amplicon Sequence Variants (ASVs). All unspecified parameters were left at their default values.

#### Metagenomic sequence data preprocessing and clustering

To preprocess the raw metagenomic samples, consisting of R1 and R2 clipped FASTQ files, we trimmed and quality-filtered the reads using the bbduk tool from the BBMap suite v38.79 (Bushnell 2014), with minimum quality and length thresholds set to 20 and 75 bp, respectively. Additionally, we employed an alternative preprocessing strategy to generate quality-checked merged paired-end reads: reads were first merged using PEAR v0.9.8 (Zhang 2014) with a minimum overlap of 12 nucleotides, then trimmed and filtered using bbduk with the same parameters.

To construct protein clusters, referred to here as Operational Protein Units (OPUs), we applied the following procedure: (1) quality-checked R1 and R2 reads were assembled into contigs using MEGAHIT v1.2.9 (Li et al. 2016) with the “meta-sensitive” preset and a minimum contig length of 250 bp; (2) quality-checked paired-end reads were mapped to the assembled contigs using the BWA-MEM algorithm from BWA v0.7.17 (Li and

Durbin 2009); (3) duplicate reads were removed with the Picard toolkit v2.24 (Broad Institute), and coverage was estimated using SAMtools v1.9 (Li 2009), considering only primary alignments with a quality score ≥10; (4) Open Reading Frames (ORFs) were predicted using FragGeneScanRs v1.1.0 (Van der Jeugt 2022), and mean coverage per ORF was calculated with Bedtools v2.30.0 (Quinlan and Hall 2010), excluding ORFs shorter than 60 bp; (5) the resulting amino acid sequences were clustered into OPUs using MMseqs2 commit version 12c499dcd647fb0d1c799bc5c6f9f25328ca3e09 (Steinegger and Söding 2018), with a minimum aligned residue fraction of 85% and identity threshold of 70%. All other parameters were set to default.

To generate the OPU abundance matrix, we summed the mean coverage of all ORFs within each OPU, excluding OPUs present in fewer than two samples or with a total abundance below 30. To account for differences in sequencing depth, the final OPU abundance matrix was rarefied to the minimum sample abundance (i.e., XXX).

**Assigning geographic coordinates to SAMO samples**

To determine the latitude and longitude coordinates to be assigned to samples from SAMO based on their similarity to six samples from the TARA Oceans miTag 16S rRNA sequences (Sunagawa 2015), we applied the following methodology: We computed the Bray Curtis Scaled Similarity (BCss), as described in Equation 1, and utilized it to compute the the weighted average of latitude and longitude coordinates, as shown in Equation 2.

Equation 1.

$$BCss_{ij} = (1 - BC_{ij})/(\sum_{j=1}^{n} 1 - BC_{ij})$$

Equation 2.

$$coordinate_i = (\sum_{j=1}^{n} \sqrt[3]{(coordinate_j)} BCss_{ij})^3$$

Where $BC_{ij}$ is the Bray–Curtis dissimilarity between sample *i* (SAMO) and *j* (TARA Oceans); $coordinate_i$ and $coordinate_j$ are the latitude (or longitude) values of samples *i* and *j*, respectively; *n* is the total number of TARA Oceans samples; $BCss_{ij}$ is the Bray–Curtis scaled similarity between samples *i* and *j*.

Note that we take the cubic root of the geographic coordinates within the summation term, and then raise the summation to the power of three in order to downweight southern samples that are more distantly located.

**Computation of OPUs and ASVs gains and losses between semesters**

To estimate ASV and OPU gains and losses between semesters, we applied the Bray–Curtis dissimilarity decomposition described in Legendre (2019), and briefly explained here. If we consider the Bray–Curtis dissimilarity as BC = (B + C)/(2A + B + C), when comparing samples i and j, the terms A, B, and C are defined as follows.

- A is the sum of all shared species between samples i and j having the lesser abundance (i.e., the sum of $A_k = \min(k_i, k_j)$, where $k_i$ and $k_j$ are the abundance of species k in sample i and j, respectively).
- B is the part of the abundance that is higher in sample i (i.e., the sum of $B_k = (k_i - k_j)$ if $k_i > k_j$; $B_k = 0$ otherwise).

- C is the part of the abundance that is higher in sample j (i.e., the sum of $C_k = (k_j - k_i)$ if $k_j > k_i$; $C_k = 0$ otherwise).

Thus, the B and C, represent the species losses and gains when going from sample i to j.

To compare gains across transitions between Summer–Fall and Winter–Spring (and vice versa), we computed the B and C components for all possible pairwise comparisons. For each transition type, we recorded the number of cases where the C component exceeded the B component and counted these as gains. Although an existing function in an R package can estimate gains and losses, it is designed for a different data structure, specifically, sample-by-sample comparisons between two separate abundance matrices. Therefore, we developed a custom implementation to perform all-vs-all comparisons within a single abundance matrix.

**Beta Nearest Taxon Index and Nearest Taxon Index**

NTI (Nearest Taxon Index) and bNTI (beta Nearest Taxon Index) are phylogenetic metrics used to study community assembly processes. NTI measures the degree of phylogenetic clustering or overdispersion within a community relative to expectations under random assembly. It is calculated as the number of standard deviations by which the observed Mean Nearest Taxon Distance (MNTD) (i.e., the mean phylogenetic distance from each ASV to its closest relative) differs from the mean MNTD of a null model distribution.

Similarly, bNTI quantifies phylogenetic turnover between two communities relative to a random expectation. It is computed as the number of standard deviations by which the observed beta Mean Nearest Taxon Distance (bMNTD) (i.e., the mean distance between each ASV in one community and its nearest phylogenetic relative in another) deviates from the mean bMNTD of a null model. Values of NTI and bNTI greater than 2 or lower than -2, indicate significantly greater or lower than expected by chance (Stegen 2012).

To compute the bNTI and NTI we applied the following methodology: 1) The 1000 most abundance ASV sequences (as suggested in Stegen 2013) were extracted from the rarefied abundance tables, and aligned against the SILVA reference alignment SILVA_138.1_SSURef_NR99_tax_silva_full_align_trunc.fasta (Quast 2013) using the PyNAST tool (Caporaso 2010). 2) The resulting alignment was quality controlled (poorly aligned regions or sequences removed) with the trimAL tool (Capella-Gutiérrez 2009) (with the parameters -gt 0.3 -st 0.001 (same as reported by Logares 2020)). 3) Aligned ASVs sequences were used to construct a phylogeny using the FastTree tool (Price 2009) (with the parameter -gtr -gamma (same as reported by Logares 2020)). 4) We The weighted bNTI was computed utilizing the code of Stegen 2013 https://github.com/stegen/Stegen_etal_ISME_2013, and weighted NTI with function 'ses.mntd' of the Picante R package version 1.8.2 (Kembel 2010). Null models were generated by 999 randomizations of the tree leafs.

Further, as required for the computation of these indices, we evaluated the presence of phylogenetic signals in ASVs optimal environmental conditions. For such analysis, we partitioned the phylogenetic distance into 50 distance bins, and computed the environmental distance between each pair samples as the euclidean distance of the scaled salinity and temperature variables. Next, we computed the optimal environmental conditions for each ASVs, as the average scaled salinity and temperature values weighted by its abundance. We then applied the Mantel test to test for significant correlations ($p < 0.05$) between phylogenetic and environmental distances within each bin. To run the Mantel test, we used the 'mantel' function from the vegan R package v2.6-8 (Oksanen 2025).

### Supplementary tables

**Supplementary table S1.** Main statistics describing the raw and preprocessed amplicon 16S rRNA dataset.

| Statistic | Raw R1 reads | Preprocessed reads |
|---|---|---|
| Total number of reads | 4,886,257 | 2,338,478 |
| Mean number of reads | 187,932.96 | 89,941.46 |
| Minimum number of reads | 6,617 | 2,693 |
| Maximum number of reads | 560,585 | 382,480 |
| Mean read length | 301 | 372.06 |

**Supplementary table S2.** Main statistics describing the metagenomic raw and preprocessed dataset.

| Statistic | Raw R1 reads | Preprocessed paired R1 reads | Preprocessed merged reads |
|---|---|---|---|
| Total number of reads | 222,226,308 | 207,252,940 | 109,837,382.5 |
| Mean number of reads | 10,101,195.82 | 9,420,588.18 | 4,992,608.3 |
| Minimum number of reads | 328,3445 | 3,064,597 | 1,624,872.5 |
| Maximum number of reads | 18502598 | 17,333,291 | 9,137,802 |
| Mean read length | 221.9 | 211.16 | 252.02 |

**Supplementary table S3.** Main statistics describing the assembled metagenomes.

| Date | N50 | L50 | N90 | L90 | Largest contig | # of contigs |
|---|---|---|---|---|---|---|
| 2018-02-20 | 503 | 66,630 | 392 | 148,808 | 7,576 | 172,847 |
| 2018-06-18 | 626 | 250,407 | 403 | 694,289 | 24,456 | 838,633 |
| 2018-11-06 | 646 | 295,628 | 403 | 846,356 | 66,046 | 1,027,050 |
| 2019-09-18 | 594 | 218,694 | 405 | 521,410 | 11,638 | 619,388 |
| 2018-11-20 | 647 | 191,061 | 403 | 560,736 | 84,576 | 681,910 |
| 2019-09-25 | 575 | 122,546 | 404 | 285,656 | 9,741 | 337,106 |

| | | | | | | |
|---|---|---|---|---|---|---|
| 2019-10-08 | 661 | 276,660 | 407 | 772,679 | 37,690 | 938,672 |
| 2020-03-04 | 738 | 314,065 | 407 | 1,037,256 | 136,485 | 1,289,249 |
| 2018-03-20 | 590 | 193,825 | 402 | 492,382 | 16,936 | 586,810 |
| 2018-12-20 | 634 | 164,448 | 408 | 437,274 | 27,788 | 525,541 |
| 2019-10-22 | 582 | 273,894 | 403 | 661,895 | 20,400 | 784,532 |
| 2020-03-11 | 599 | 261,913 | 397 | 720,794 | 59,208 | 864,938 |
| 2018-04-09 | 571 | 61,580 | 402 | 147,859 | 6,992 | 174,631 |
| 2019-11-06 | 620 | 167,282 | 406 | 430,098 | 3,975 | 514,725 |
| 2018-04-18 | 594 | 163,625 | 402 | 418,867 | 101,361 | 499,698 |
| 2019-07-11 | 604 | 265,596 | 396 | 747,257 | 50,904 | 900,518 |
| 2019-11-21 | 652 | 264,734 | 411 | 699,898 | 15,551 | 845,341 |
| 2018-04-25 | 650 | 85,616 | 406 | 238,223 | 20,843 | 289,513 |
| 2019-08-28 | 666 | 169,113 | 399 | 558,884 | 296,760 | 686,842 |
| 2019-12-04 | 710 | 214,952 | 407 | 668,786 | 91,925 | 824,480 |
| 2018-10-24 | 636 | 236,760 | 407 | 623,031 | 37,730 | 748,841 |
| 2019-12-17 | 697 | 292,572 | 405 | 888,415 | 73,979 | 1,095,096 |

**Supplementary table S4.** Latitude and longitude of the six TARA Oceans samples compared against SAMO samples.

| Sample | Latitude | Longitude | Relative to SAMO |
|---|---|---|---|
| TARA_072_SRF_0.22-3 | -8.7789 | -17.9099 | North |
| TARA_076_SRF_0.22-3 | -20.9354 | -35.1803 | North |
| TARA_078_SRF_0.22-3 | -30.1367 | -43.2899 | North |
| TARA_082_SRF_0.22-3 | -47.1863 | -58.2902 | South |
| TARA_084_SRF_0.22-3 | -60.2287 | -60.6476 | South |
| TARA_085_SRF_0.22-3 | -62.0385 | -49.529 | South |

**Supplementary table S5.** Detailed statistics of the correlations between axes of the NMDS analyses and the environmental variables temperature and salinity

| Type of data | MDS1 vs Temperature | MDS2 vs Salinity |
|---|---|---|
| ASV | Pearson's r = -0.78<br>P-value = 2.6e-06 | Pearson's r = 0.776<br>P-value = 3.1e-06 |
| OPU | Pearson's r = -0.896<br>P-value = 1.667e-08 | Pearson's r = 0.725<br>P-value = 1.3e-4 |

### Supplementary figures

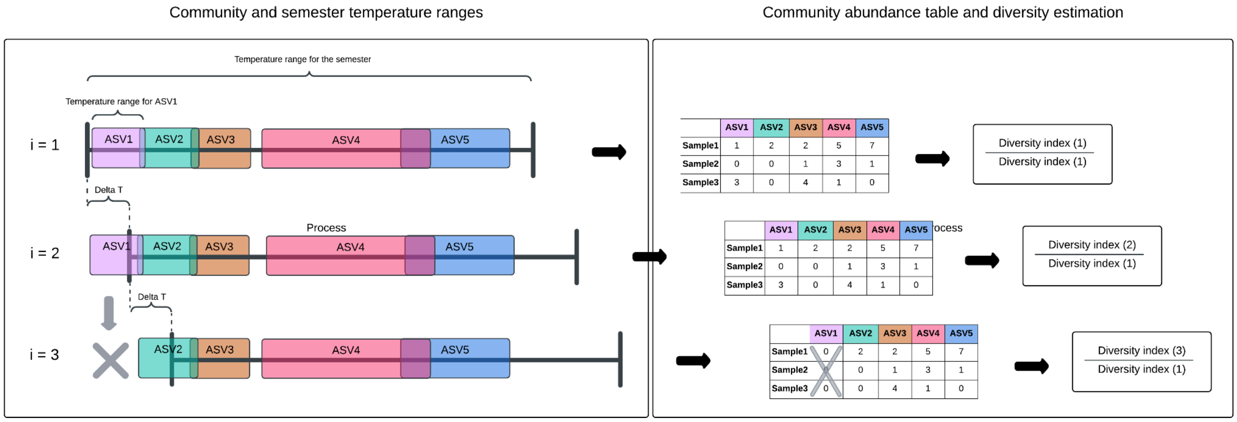


**Supplementary figure S1. Schematic representation of the methodology used to simulate increased temperature scenarios.** For illustration, only three simulation steps are shown, involving five ASVs. The left panel depicts the community and semester temperature ranges under the simulated temperature increase. Colored boxes represent the temperature tolerance ranges of each ASV, while the black whiskers show the temperature range for the target semester (either Summer–Fall or Winter–Spring). In each iteration, the semester's temperature range is artificially shifted upward in increments of 0.2 °C (i.e., delta T), up to a total of 3 °C. ASVs whose maximum temperature falls below the new minimum of the adjusted range are assumed to be excluded from the community and are assigned an abundance of zero. The right panel shows the resulting abundance matrices for each iteration and the corresponding calculation of diversity indices. Diversity is assessed using the Shannon index and richness (observed and Chao1), scaled relative to the original community before any temperature modification. In the example, from iteration 1 to 2, all ASVs remain within the temperature range, and the community composition, and thus diversity, remains unchanged. However, in iteration 3, ASV1's temperature range no longer overlaps with the shifted semester range, leading to its exclusion from the community prior to diversity calculation.

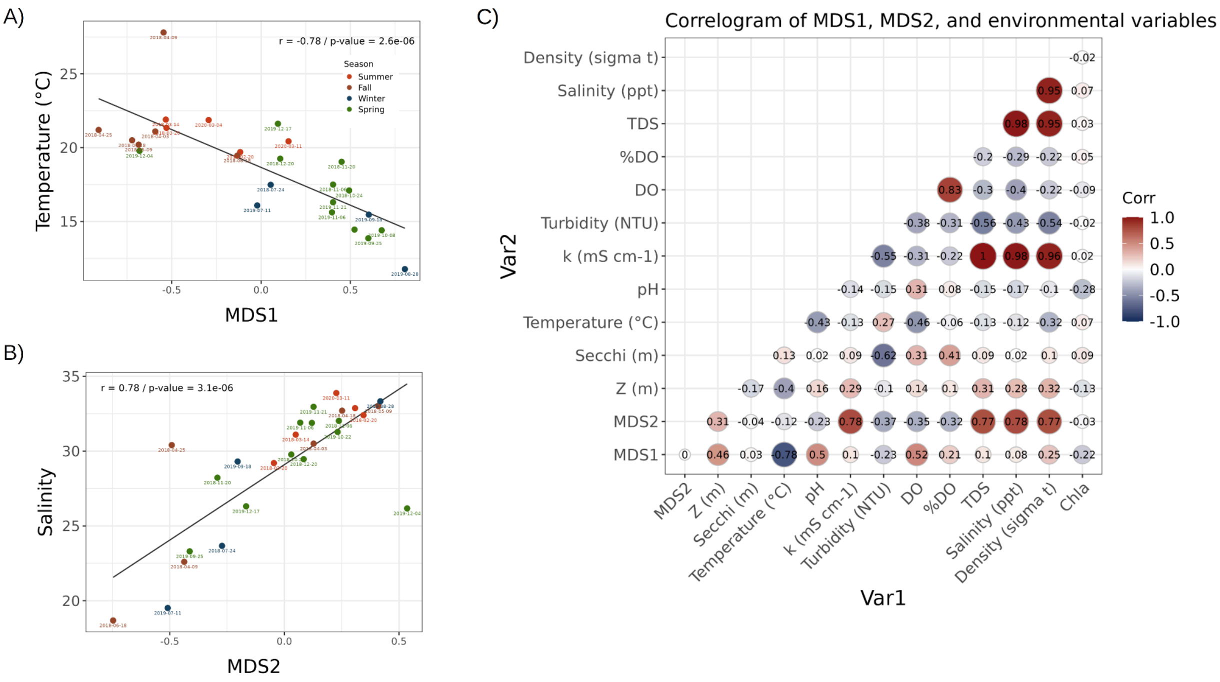


**Supplementary figure S2. Correlation of environmental variables with MDS1 and MDS2 axes from the NMDS based on ASV Bray–Curtis dissimilarities.** A–B) Scatter plots showing strong negative correlations between MDS1 and MDS2 with temperature and salinity, respectively. Points are colored by season. C) Correlogram displaying Pearson correlation coefficients between MDS axes (MDS1, MDS2) and environmental variables. Circle size and color indicate the strength and direction of the correlations, with red representing positive and blue representing negative relationships. Correlation values are shown within each cell.

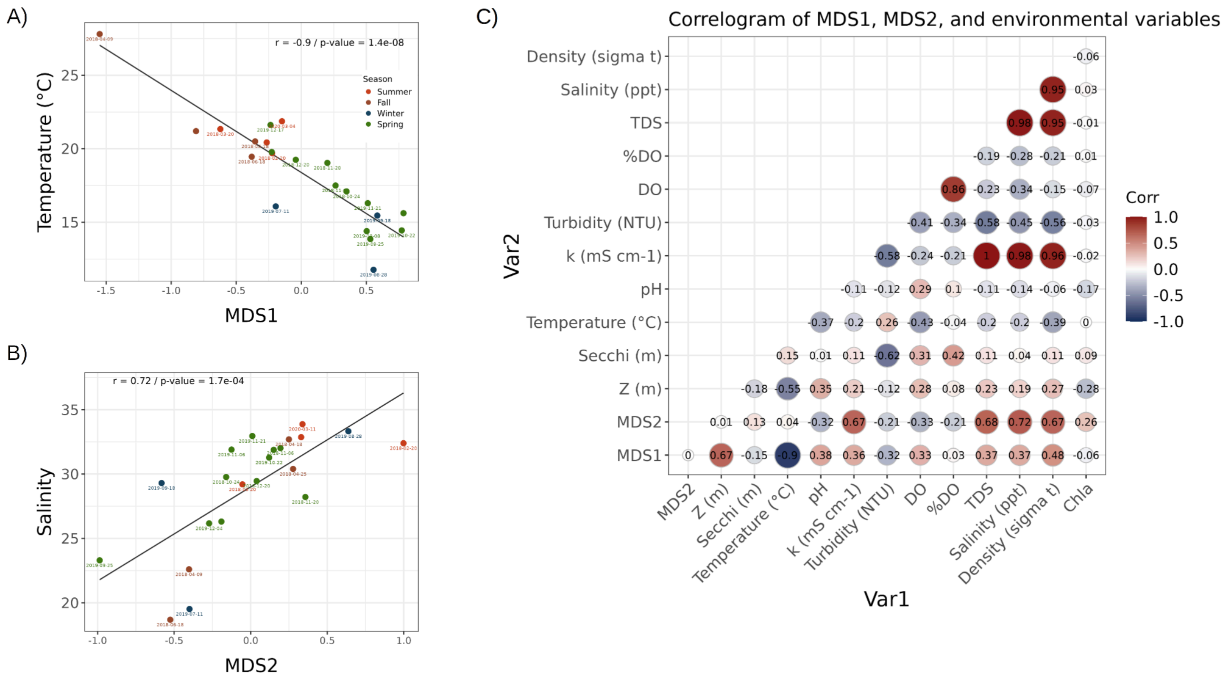


**Supplementary figure S3. Correlation of environmental variables with MDS1 and MDS2 axes from the NMDS based on OPU Bray–Curtis dissimilarities.** Same plots as in Supplementary Figure S2. A–B) Scatter plots showing correlations between NMDS axes (MDS1, MDS2) and temperature and salinity. C) Correlogram summarizing Pearson correlations between MDS axes, environmental variables.

A)

B)

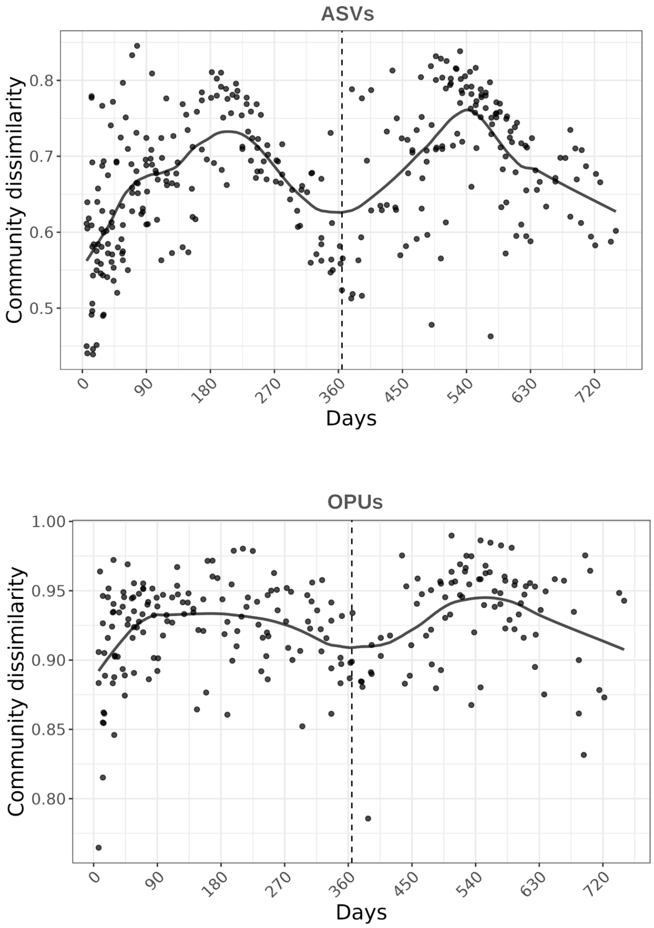


**Supplementary figure S4. Time-decay relationship over a two-year period.** A) and B) Bray–Curtis dissimilarity between sample pairs, based on rarefied ASV and OPU compositions, respectively, plotted against the time difference between samples (in days). LOESS regression curves are shown to illustrate temporal trends in community dissimilarity. The vertical dashed line at day 365 marks sample pairs separated by approximately one year.

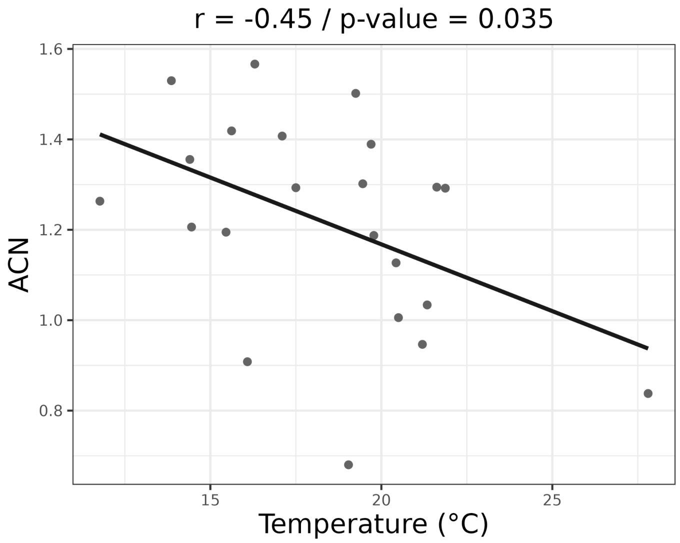


**Supplementary figure S5. Relationship between average 16S rRNA gene copy number (ACN) and temperature.** The scatter plot shows a significant negative Pearson correlation between temperature and ACN across samples (Pearson's $r$ = –0.45, $p$ = 0.035). The solid black line represents the linear regression fit. ACN was estimated from merged, assembled metagenomic samples using the ACN tool (Pereira-Flores 2019).

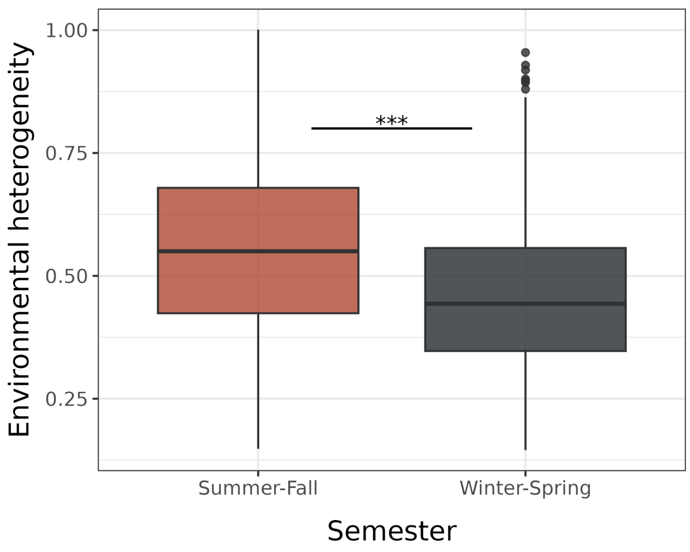


**Supplementary figure S6. Environmental heterogeneity across seasons (semesters) in coastal marine environments.** Boxplot comparing the distribution of environmental heterogeneity values between the Summer–Fall and Winter–Spring semesters. Each box represents the interquartile range (IQR), with the horizontal line indicating the median and whiskers extending to 1.5 times the IQR; outliers are shown as individual points. Environmental heterogeneity was estimated based on scaled Euclidean distances among samples using temperature, salinity, pH, chlorophyll-a, and dissolved oxygen. Results indicate significantly greater heterogeneity in the Summer–Fall period compared to Winter–Spring (Welch's ANOVA, $p = 2.506e{-}05$).

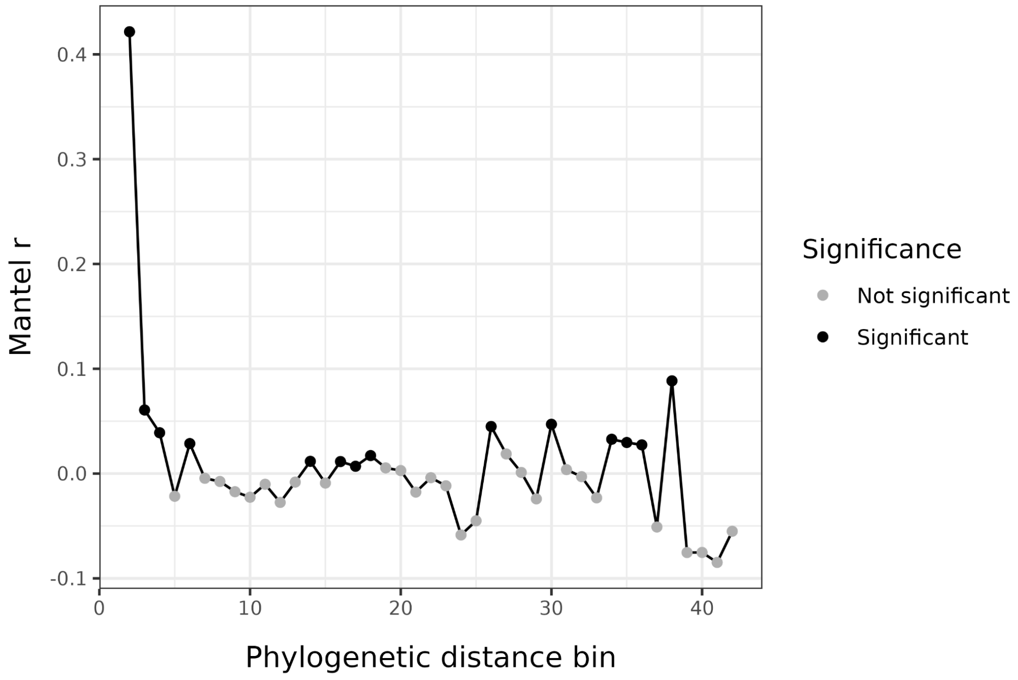


**Supplementary figure S7. Test for phylogenetic signal over short phylogenetic distances.** The plot shows Mantel correlation coefficients (*r*) from tests comparing environmental and phylogenetic distances across increasing phylogenetic distance bins (x-axis). Black and gray points indicate bins with significant and non-significant correlations ($p < 0.05$), respectively. A strong and significant positive correlation in the first bin suggests that environmental similarity is more conserved among closely related taxa, supporting the assumption that ecological traits influencing environmental associations exhibit phylogenetic signals at short evolutionary distances (Stegen 2012).

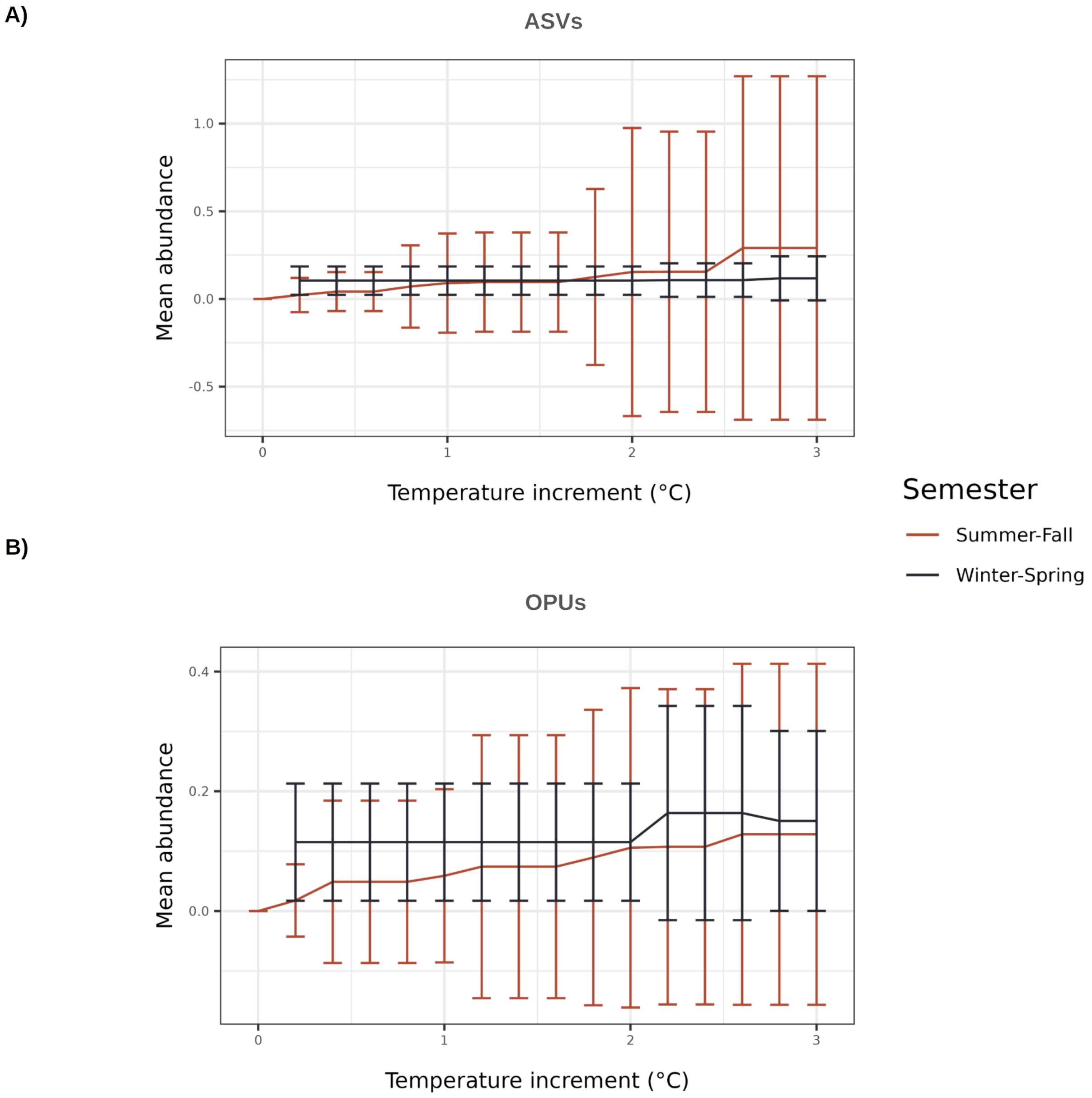


**Supplementary Figure S8. Impact of simulated temperature increases on mean microbial abundance across seasons.** A–B) Changes in mean abundance of ASVs and OPUs, respectively, under simulated temperature increases from 0 °C to 3 °C. Summer–Fall and Winter–Spring semesters are represented by red and dark blue lines, respectively. Error bars indicate standard deviation across samples. No significant differences between semesters were observed in the mean abundance of ASVs or OPUs lost due to temperature shifts (Welch's ANOVA, $p > 0.1$).
